# The SARS-CoV-2 cellular receptor ACE2 is expressed in oropharyngeal cells and is modulated in vitro by the bacterial lysate Lantigen B

**DOI:** 10.1101/2022.05.06.490962

**Authors:** Caterina Pizzimenti, Antonella D’Agostino, Paola Pirrello, Alessia Ruiba, Giovanni Melioli

## Abstract

Angiotensin-converting enzyme2 (ACE2) is the main cell surface receptor of the SARS-CoV-2 spike protein and is expressed in a variety of cell types, including cells of the respiratory tract. A bacterial lysate used for the prophylaxis of respiratory infections (OM-85), was recently shown to downregulate the expression of ACE2 in epithelial cells, suggesting its possible role as a prophylaxis of the onset of COVID19. Another bacterial lysate (Lantigen B, administered sublingually) is used in the prophylaxis of recurrent respiratory tract infections. It contains antigens obtained by chemical lysis from the most representative microbes of the respiratory tract. In this in vitro study, the capacity of Lantigen B to decrease ACE2 in human oropharyngeal cells was evaluated. The study was carried out in 40 healthy donors undergoing oropharyngeal swab for routine SARS-CoV-2 detection. Cells were treated in vitro with a 1:2 of Lantigen B. ACE2 expression was evaluated using a fluorescent anti-ACE2 monoclonal antibody and flow cytometry. A reduction in the number of positive cells was observed in 72% of the patients, while a modulation of ACE2 expression was observed in 62% of the samples. As a control, the expression of the CD54 rhinovirus receptor in the same cells was unaffected. To evaluate the functional effects of down regulation, in a subset of samples, the same oropharynx cells were incubated with Lantigen B and infected with wild-type SARS-CoV-2. After 24 hours, viral RNA, as assessed by rt-PCR, was significantly lower in samples treated with Lantigen B. In conclusion, this study demonstrates that Lantigen B, at a pharmacological dose, modulates the expression of the main SARS-CoV-2 receptor in oropharyngeal cells, and reduces viral yield. This activity could be synergistic with other approaches (vaccination and therapy) by reducing the number of potentially infected cells and thus reducing the effects of SARS-CoV-2 infection.

## Introduction

The COVID-19 pandemic strongly stimulated basic and applied research. In a few weeks, real-time polymerase chain reaction tests for virus identification were developed and distributed worldwide. Panels of humanized monoclonal antibodies have been obtained [1] and the use of these tools has begun in clinics [2]. Then active vaccines have been developed and introduced into humans [3, 4] and, at present, it has been calculated that almost 64% of the entire human population has been vaccinated (https://ourworldindata.org/covid-vaccinations?country=OWID_WRL). Finally, novel drugs have been proposed or are being developed [5]. The SARS-CoV-2 variants appeared early and, at present, the family of Omicron variants is widely distributed. Despite the use of all these tools, COVID-19 remains a relevant cause of death in elderly and at risk populations.

The cell surface molecule Angiotensin Converting Enzyme 2 (ACE2) is the main receptor for the attachment of the SARS-CoV-2 spike protein (S) [6,7]. Together with other mechanisms, binding of the spike protein to the ACE2 receptor allows the fusion of the virus and the cell membrane, thus allowing the infection of target cells. ACE2 has been detected in many different cell types, including nasal, intestinal, kidney, testes, gallbladder, and heart cells, and, to a lesser extent, tongue, tonsil, and olfactory region cells [8,9]. Therefore, oropharynx cells represent the first target of SARS-CoV-2, and any attempt to increase the resistance of these cells to SARS-CoV-2 infection seems to be interesting.

In a recent article [10], D. Vercelli and co-workers have shown that a bacterial lysate developed decades ago, OM-85, has the ability to reduce the expression of ACE2, both in cell lines and primary cells of human airways. On these bases, the authors suggest that the administration of OM-85, which reduces ACE2 expression, could to some extent inhibit respiratory tract target cell infection. Thus, OM-85 could act as a prophylactic treatment that can be used in association with vaccines and other future therapies to control the spread of COVID-19.

Bacterial lysates were developed more than 40 years ago and have been shown to reduce the number and severity of respiratory tract infections (or exacerbations) in treated patients [11,12]. Details of the mechanisms are not fully described, although the capacity to activate dendritic cells, inducing a specific and functionally effective immune response, and recruiting innate immune-competent cells has been observed in the past [13,14]. Furthermore, epithelial cells have been shown to be able to interact with certain bacterial lysates (such as polyvalent mechanical bacterial lysates), resulting in epithelial cell activation, proliferation, and tight junction sealing [15]. Lantigen B is a prototype of bacterial lysates: it was originally developed in the 1960s, and since its availability, it has been empirically used in reducing the number of respiratory tract infections during the winter season. Lantigen B contains the particulate and soluble fraction obtained by chemical lysis of six different microbes of the respiratory tract (*Staphylococcus aureus*, *Klebsiella pneumoniae*, *Streptococcus pneumoniae*, *Streptococcus pyogenes*, *Moraxella catarrhalis*, and *Haemophilus influenzae*). Several articles described this feature: in particular, a recent double blind placebo-controlled clinical trial has further demonstrated its clinical activity [16]. Lantigen B is administered under the tongue; thus, the drug enters into direct contact with the epithelial cells of the mouth and the oropharynx. If modulation of ACE2 expression is a characteristic of bacterial lysates (and not only of OM-85), a possible effect of Lantigen B on the same receptor structure could be hypothesized. In this report, we show that oropharynx epithelial cells express ACE2 and that Lantigen B is capable of reducing in vitro the expression of ACE2 after 24-hour treatment at doses comparable to those used in vivo.

## Materials and Methods

During a period of a few weeks from February to April 2022, epithelial cells were collected during routine pharyngeal swab for SARS-CoV-2 detection from a group of researchers and employees (14 men and 16 women, age ranging from 35 to 73) at Bruschettini Ltd, the Italian firm that produces Lantigen B. All donors gave a verbal informed consent for the use of leftover cells after sampling and rt-PCR analysis. All donors were negative for SARS-CoV-2 antigens and RNA. All donors leftover cells were therefore used for the study. Cells were washed in saline and then resuspended in RPMI 1640 (Euroclone, Pero, Italy) with 10% foetal bovine serum (Euroclone, Pero, Italy) and gentamycin (160 mg/L). Cell viability was scored by Trypan blue exclusion.

Before treating oropharyngeal cells with Lantigen B, the presence of ACE2 on the surface of oropharyngeal cells was evaluated in freshly collected cells from 15 donors, using a specific anti-ACE2 monoclonal antibody (R&D Systems, labelled Alexa Fluor 488) and flow cytometry (Cytoflex, Beckman Coulter). Briefly, a proper dilution of anti-ACE2 monoclonal antibody (mAb) was added to cells and allowed to react at 4 °C for 30 minutes. Cells were analysed by flow cytometry. An isotype control (FITC-labelled anti-CD83, Beckman Coulter) was used to evaluate possible nonspecific binding to epithelial cells. The results were reported as percentage of positive cells and median surface antigen.

Modulation of ACE2 by Lantigen B was evaluated in donor oropharyngeal cells in vitro under different experimental conditions. First, cells were incubated overnight in RPMI with 10% foetal bovine serum (HyClone, Sigma, USA) and gentamycin in the absence of any stimulus, as an unstimulated control. Second, the cells were incubated with the diluent of Lantigen B bacterial antigens, which contained water, polysorbate 80, chlorhexidine diacetate and sodium methyl parahydroxybenzoate. Taking into account the hypotonicity of water, a proper dilution of a 10x phosphate buffered saline (PBS) solution was added. This condition was considered the experimental negative control. Third, cells were diluted 1:2 with the drug at the dose used in humans (final 3.5 mg/mL of bacterial lysate) after reconstitution of isotonicity with 10x PBS. The drug contains antigens of the above-mentioned microbes of the respiratory tract. After 24 hours of culture, cells were collected and viability was evaluated using trypan blue dye exclusion.

The expression of ACE2 in epithelial cells was measured by the specific anti-ACE2 mAb and flow cytometry. The isotype control (FITC-labelled anti-CD83, Beckman Coulter) was also used to rule out nonspecific binding. Furthermore, to evaluate whether modulation of ACE2 can be considered in the context of a more general mechanism, another surface molecule (CD54 – ICAM-1, receptor for rhinoviruses) [16] expression modulation was investigated after Lantigen B treatment. Finally, 6 μm polystyrene microspheres (Polysciences, USA) were used to calibrate the flow cytometer and evaluate cell size. The ACE2 modulation was performed in three different sessions using three different cohorts consisting of 15, 13 and 12 donors, respectively. All tests were performed in duplicates. To evaluate the functional effects of ACE2 downregulation of oropharyngeal cells, in four selected samples, the capacity of Lantigen B to reduce the virus yield was evaluated, Briefly, 0.05 mL from transport medium of three different positive samples was added to 0.1 mL of oropharyngeal cells in RPMI 1640 as previously described before and to 0.05 mL of either saline or Lantigen B in sealed microtubes. After 24 hours, virus RNA was extracted from the cell supernatants at 98°C for 3 minutes. The extracts were assayed by rt-PCR (COVID-19 screening, Clonit, Italy) according to the producer’s instructions.

## Results

### Expression of ACE2 in normal oropharyngeal cells

As described, the first series of experiments focused on identifying ACE2 in normal oropharyngeal cells immediately after cell collection with an oropharyngeal swab. In fact, only in the presence of clear ACE2 expression could any attempt to down-regulate its expression be evaluated. Figure 1 shows the expression of ACE2 in the FITC channel and the same population treated with an isotype control. As described, the assay was performed on cells from 15 different donors. ACE2 was expressed on the surface of oropharyngeal cells with some heterogeneity. The median fraction of ACE2-positive cells was 68% (ranging from 52 to 80%). Few samples (2 out of 15) were characterized by a reduced percentage of ACE2-positive cells (ranging from 37 to 47%). The intensity of fluorescence was more homogeneous. In particular, the epithelial cells were large; the diameter measured using 6 μm plastic beads in the forward scatter channel ranged between 18 and 56 μm. Under these conditions, the very large surface area caused a strong signal in the FITC channel. Furthermore, autofluorescence was high but significantly lower than that of cells stained with ACE2 mAb.

**Figure 1.**
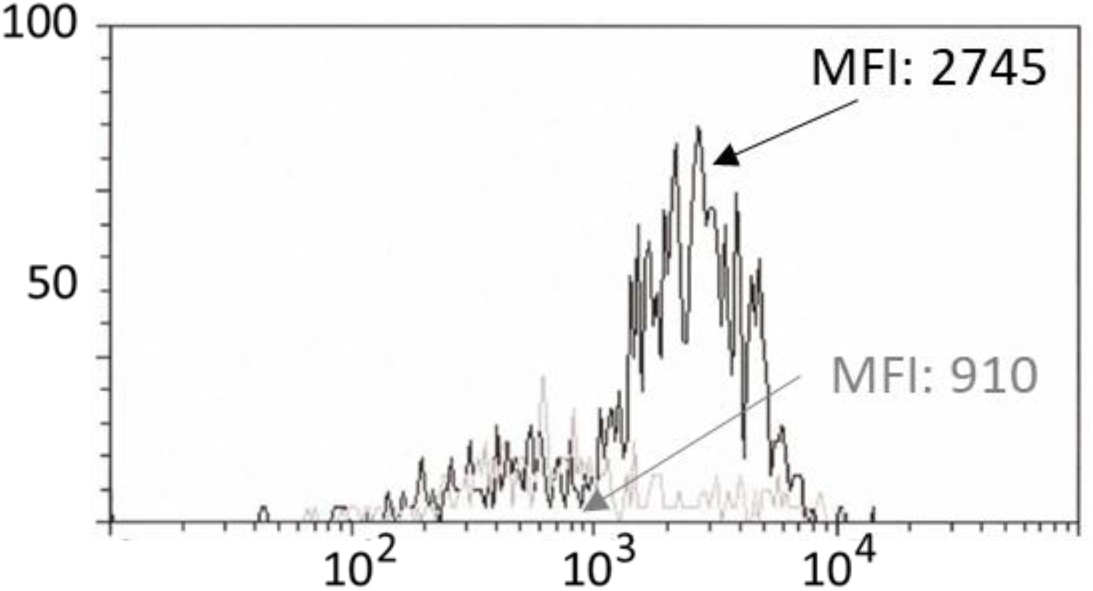
Demonstration of the presence of the ACE2 protein on the surface of oropharyngeal cells. The black profile represents the distribution of ACE2 on the cell surface. In gray, the isotypic negative control shows weak autofluorescence.

### Modulation of ACE2 in normal oropharyngeal cells after incubation with Lantigen B

Using the same conditions as in the first experiments, donor oropharyngeal cells were tested for the presence of ACE2 after 24 hours of incubation in the presence and absence of Lantigen B. Treatment with Lantigen B resulted in a reduction in the number of cells in 72% of samples. The median reduction in the number of positive cells was 44%, ranging from 18% to 69.5% in the group of 40 donors tested in three different sessions. A reduction in fluorescence intensity (representing the number of surface molecules) was observed in 26 samples (65%, median reduction 12.4%, ranging from 4.2 to 43%) under the same conditions. Figure 2 shows the results of the two different donors. Notably, not all samples where the percentage of ACE2-positive cells was reduced also showed a reduction in fluorescence intensity.

**Figure 2.**
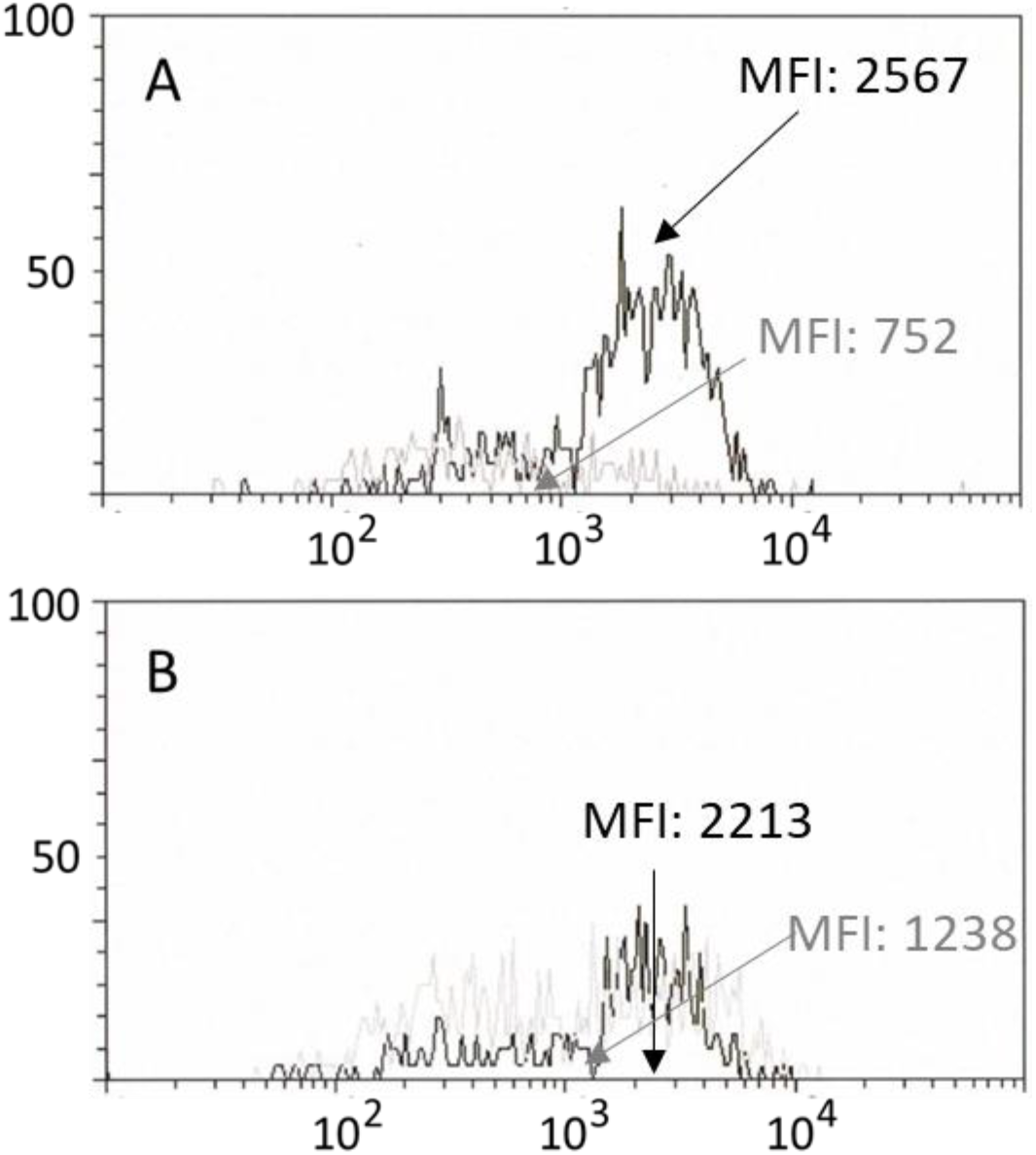
Down-regulation of ACE2 in oropharyngeal cells. Analyses were performed on oropharyngeal cells after 24-hour incubation in the presence of pharmacological doses of Lantigen B. In black, cells from donors treated with drug excipients alone. In gray, cells from donors after drug treatment. Panel A shows a representative example of the down-regulation observed in 72% of donor samples. In panel B, a different behaviour (absent or poor reduction in ACE2 expression) was observed in the remaining samples.

CD54 (ICAM-1) was used as a control, being the receptor of the rhinovirus family. Although well expressed on the surface of almost all oropharyngeal cells, its intensity of expression, evaluated in the last group of 12 donors, as well as the number of positive cells, remained unchanged with the administration of Lantigen B in this experimental setting.

The effect of Lantigen B on virus proliferation was tested using a non-conventional test as described. The results of representative experiments are shown in Figure 3. All samples treated with Lantigen B reduced the virus yield measured as RNA in rt-PCR. The median reduction was 9.2 times (ranging from 1.4 to 45 times).

**Figure 3.**
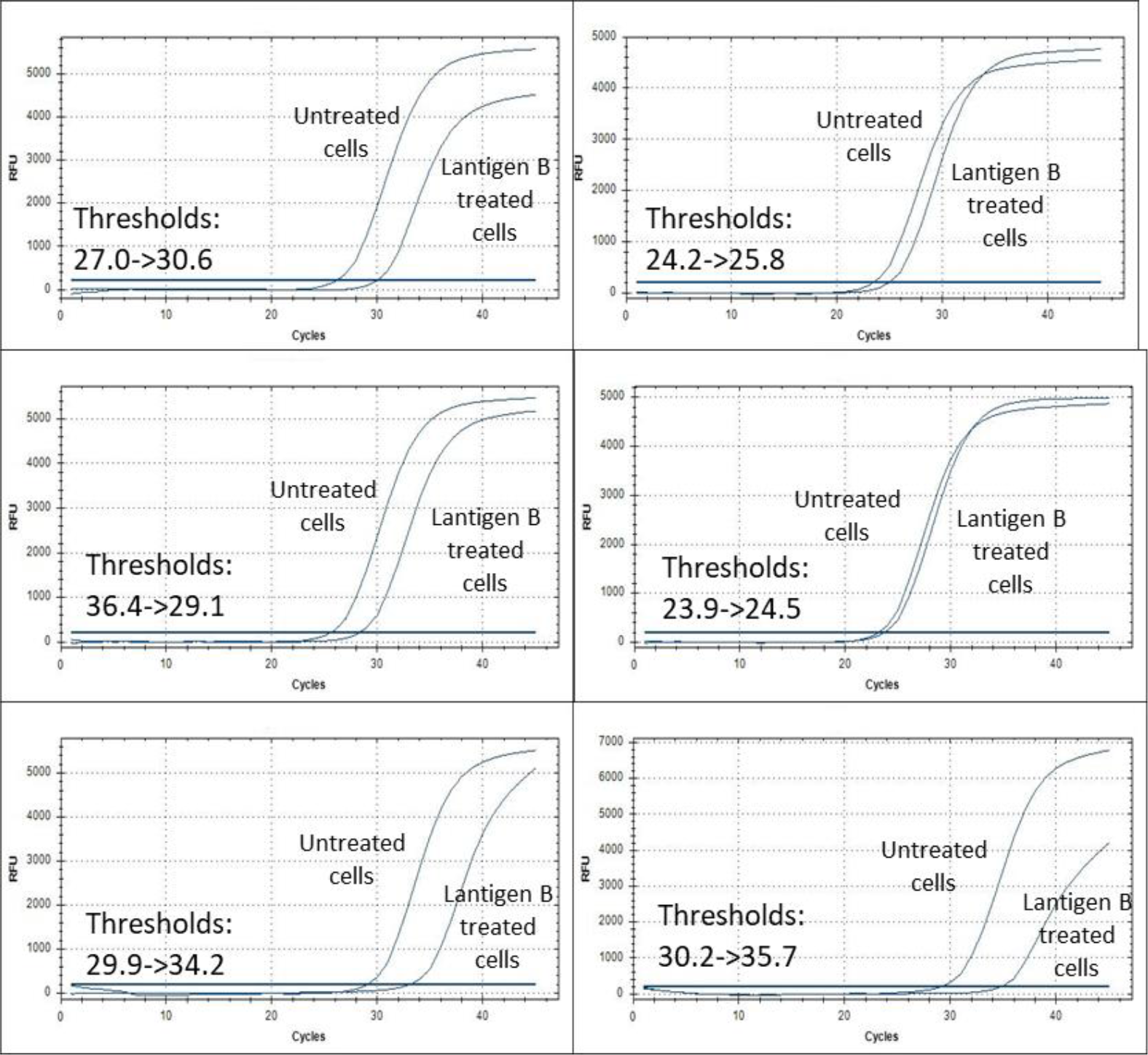
Decrease of SARS-CoV-2 RNA copies upon 24h Lantigen B treatment. Graphs show SARS-CoV-2 RNA detection via rt-PCR from donors’ oropharynx cells.

## Discussion

A recent article by Vercelli and co-workers showed that a bacterial lysate (OM-85) has the ability to reduce the expression of ACE2 in epithelial cells after in vitro culture [10]. In the same experiments, OM-85 was shown to inhibit infection of epithelial cells. This result is interesting because it can be hypothesized that drugs with this capability can be used in the context of COVID-19 prophylaxis. In this context, vaccination has been shown to induce a strong systemic immunological response that lasts for months [17]. However, in the same patients, the locoregional response at the mucosal level is shorter [18,19]. These findings could be one of the explanations for the inability of vaccination to prevent reinfections, whereas severe disease is better controlled by systemic immunity. Thus, already available drugs that can reduce cell infection, representing the first barrier, are extremely useful. However, these results on M-85 were obtained using high concentrations of bacterial lysate (from 0.24 to 1.92 mg/mL), doses that seem to be difficult to reach in vivo, considering that a single pill of OM-85 contains 7 mg of active principle.

Lantigen B is administered at the mucosal level, and this could be an added value of this treatment. In fact, in vitro tests on freshly isolated oropharyngeal epithelial cells indicate that a significant reduction in ACE2 expression is achieved using drug concentrations that are easily reached in vivo. Additionally, ACE2 expression reduction is noticed after 24 hours, while OM-85 has effects observed after a longer period under the experimental conditions described.

This effect appears to be strictly related to the presence of bacterial antigens, as the excipients and preservatives do not have an effect when administered alone. Furthermore, the surface expression of CD54, a rhinovirus receptor, was not modified by in vitro treatment with Lantigen B, suggesting that, at least under these experimental conditions, down-regulation of ACE2 seems to related to a specific mechanism.

One apparent limitation of this study is represented by the absence of a standard assay on Lantigen B-treated, SARS-CoV-2 infected oropharyngeal cells. However, a non-conventional test was performed by adding normal oropharyngeal cells to the transport medium of positive SARS-CoV-2 samples, either in the presence or in the absence of Lantigen B. The median reduction in viral RNA was 9.2 fold and this finding appears to be strongly related to the downregulation of ACE2 expression mediated by Lantigen B in target cells. Notably, this 10-fold reduction of virus yield, evaluated by rt-PCR is similar to the results observed by Vercelli and co-workers using the reduction of the number of plaque forming units [10].

## Conclusions

Lantigen B, a bacterial lysate used for decades in the prophylaxis of recurrent respiratory infections, has a well-established activity in reducing the number of infections in clinical trials. Our novel findings add that Lantigen B has activities that could be useful in the prophylaxis of COVID-19 (or at least in the prevention of SARS-CoV-2 asymptomatic infections), particularly in the period that seems to be less protected by vaccines. Thus, in addition to the well-known effects of bacterial lysates on the adaptive and innate immune response, the finding of this capability of Lantigen B adds to the mechanism of action of bacterial lysates.

## Data availability

The raw data were stored in an MS Excel file and are available upon request.

## Acknowledgements

The assistance of Alessandra Cavalieri in the preparation of the manuscript was appreciated.

## References

1. Zost, S.J., Gilchuk, P., Chen, R.E. et al. Rapid isolation and profiling of a diverse panel of human monoclonal antibodies targeting the SARS-CoV-2 spike protein. Nat Med 26, 1422–1427 (2020). https://doi.org/10.1038/s41591-020-0998-x

2. Hwang, YC., Lu, RM., Su, SC. et al. Monoclonal antibodies for COVID-19 therapy and SARS-CoV-2 detection. J Biomed Sci 29, 1 (2022). https://doi.org/10.1186/s12929-021-00784-w

3. Mulligan MJ, Lyke KE, Kitchin N, Absalon J, Gurtman A, Lockhart S, et al. Phase I/II study of COVID-19 RNA vaccine BNT162b1 in adults. Nature. 2020;586:589–93.

4. Wang Z, Schmidt F, Weisblum Y, Muecksch F, Barnes CO, Finkin S, et al. mRNA vaccine-elicited antibodies to SARS-CoV-2 and circulating variants. Nature. 2021;592:616–22

5. Zhao L, Li S, Zhong W. Mechanism of action of small-molecule agents in ongoing clinical trials for SARS-CoV-2: a review. Front Pharmacol. 2022;13:840639.

6. Zhu N, Zhang D, Wang W, Li X, Yang B, Song J, et al. A novel coronavirus from patients with pneumonia in China, 2019. N Engl J Med. 2020;382:727–33.

7. Hoffmann M, Kleine-Weber H, Schroeder S, Krüger N, Herrler T, Erichsen S, et al. SARS-CoV-2 cell entry depends on ACE2 and TMPRSS2 and is blocked by a clinically proven protease inhibitor. Cell. 2020;181:271–80.e8.

8. Hikmet F, Méar L, Edvinsson Å, Micke P, Uhlén M, Lindskog C. The protein expression profile of ACE2 in human tissues. Mol Syst Biol. 2020;16:e9610.

9. Hamming I, Timens W, Bulthuis ML, Lely AT, Navis G, van Goor H. Tissue distribution of ACE2 protein, the functional receptor for SARS coronavirus. A first step in understanding SARS pathogenesis. J Pathol. 2004;203:631–7.

10. Pivniouk V, Pivniouk O, DeVries A, Uhrlaub JL, Michael A, Pivniouk D, et al. The OM-85 bacterial lysate inhibits SARS-CoV-2 infection of epithelial cells by downregulating SARS-CoV-2 receptor expression. J Allergy Clin Immunol. 2022;149:923–33.e6.

11. Braido F, Tarantini F, Ghiglione V, Melioli G, Canonica GW. Bacterial lysate in the prevention of acute exacerbation of COPD and in respiratory recurrent infections. Int J Chron Obstruct Pulmon Dis. 2007;2:335–45.

12. Villa E, Garelli V, Braido F, Melioli G, Canonica GW. May we strengthen the human natural defenses with bacterial lysates? World Allergy Organ J. 2010;3:S17–23.

13. Ricci R, Palmero C, Bazurro G, Riccio AM, Garelli V, Di Marco E, et al. The administration of a polyvalent mechanical bacterial lysate in elderly patients with COPD results in serological signs of an efficient immune response associated with a reduced number of acute episodes. Pulm Pharmacol Ther. 2014;27:109–13.

14. Lanzilli G, Traggiai E, Braido F, Garelli V, Folli C, Chiappori A, et al. Administration of a polyvalent mechanical bacterial lysate to elderly patients with COPD: effects on circulating T, B and NK cells. Immunol Lett. 2013;149:62–7.

15. Migliore GS, Campana S, Pasquale CD, Carrega P, Ferlazzo G: Human airway epithelial cells directly recognize mechanical bacterial lysates eliciting tight junction sealing, antimicrobial peptides production and epithelial cell proliferation, vol. 20. Dubai, UAE: XII World Congress on COPD, Asthma & Respiratory Allergy; 2018.

16. Braido F, Melioli G, Candoli P, Cavalot A, Di Gioacchino M, Ferrero V, et al. The bacterial lysate Lantigen B reduces the number of acute episodes in patients with recurrent infections of the respiratory tract: the results of a double blind, placebo controlled, multicenter clinical trial. Immunol Lett. 2014;162:185–93.

17. Zhang Z, Mateus J, Coelho CH, Dan JM, Moderbacher CR, Gálvez RI, Cortes FH, Grifoni A, Tarke A, Chang J, Escarrega EA, Kim C, Goodwin B, Bloom NI, Frazier A, Weiskopf D, Sette A, Crotty S. Humoral and cellular immune memory to four COVID-19 vaccines. Cell. 2022 Jul 7;185(14):2434–2451.e17. doi: 10.1016/j.cell.2022.05.022. Epub 2022 May 27. PMID: 35764089; PMCID: PMC9135677.

18. Azzi L, Dalla Gasperina D, Veronesi G, Shallak M, Ietto G, Iovino D, et al. Mucosal immune response in BNT162b2 COVID-19 vaccine recipients. EBioMedicine. 2022;75:103788.

19. Sheikh-Mohamed S, Isho B, Chao GYC, Zuo M, Cohen C, Lustig Y, et al. Systemic and mucosal IgA responses are variably induced in response to SARS-CoV-2 mRNA vaccination and are associated with protection against subsequent infection. Mucosal Immunol. 2022:1–10. Doi: 10.1038/s41385-022-00511-0

